# Evolution of irreversible somatic differentiation

**DOI:** 10.1101/2021.01.18.427219

**Authors:** Yuanxiao Gao, Hye Jin Park, Arne Traulsen, Yuriy Pichugin

## Abstract

A key innovation emerging in complex animals is irreversible somatic differentiation: daughters of a vegetative cell perform a vegetative function as well, thus, forming a somatic lineage that can no longer be directly involved in reproduction. Primitive species use a different strategy: vegetative and reproductive tasks are separated in time rather than in space. Starting from such a strategy, how is it possible to evolve life forms which use some of their cells exclusively for vegetative functions? Here, we developed an evolutionary model of development of a simple multicellular organism and found that three components are necessary for the evolution of irreversible somatic differentiation: (i) costly cell differentiation, (ii) vegetative cells that significantly improve the organism’s performance even if present in small numbers, and (iii) large enough organism size. Our findings demonstrate how an egalitarian development typical for loose cell colonies can evolve into germ-soma differentiation dominating metazoans.

## 1 Introduction

In complex multicellular organisms, different cells specialise to execute different functions. These functions can be generally classified into two kinds: reproductive and vegetative. Cells performing reproductive functions contribute to the next generation of organisms, while cells performing vegetative function contribute to sustaining the organism itself. In unicellular species and simple multicellular colonies, these two kinds of functions are performed at different times by the same cells – specialization is temporal. In more complex multicellular organisms, specialization transforms from temporal to spatial Mikhailov et al. [2009], where groups of cells focused on different tasks emerge in the course of organism development.

Typically, cell functions are changed via differentiation, such that a daughter cell performs a different function than the maternal cell. The vast majority of metazoans feature a very specific and extreme pattern of cell differentiation: any cell performing vegetative functions forms a somatic lineage, i.e. producing cells performing the same vegetative function – somatic differentiation is irreversible. Since such somatic cells cannot give rise to reproductive cells, somatic cells do not have a chance to pass their offspring to the next generation of organisms. Such a mode of organism development opened a way for deeper specialization of somatic cells and consequently to the astonishing complexity of multicellular metazoans. In *Volvocales* – a group of green algae serving as a model species for evolution of multicellularity – the emergence of irreversibly differentiated somatic cells is the hallmark innovation marking the transition from colonial life forms to multicellular species Kirk [2005].

While the production of individual cells specialized in vegetative functions comes with a number of benefits Grosberg and Strathmann [2007], the development of a dedicated vegetative cell lineage that is lost for organism reproduction is not obviously a beneficial adaptation. From the perspective of a cell in an organism, the guaranteed termination of its lineage seems the worst possible evolutionary outcome for itself. From the perspective of entire organism, the death of somatic cell at the end of the life cycle is a waste of resources, as these cells could in principle become parts of the next generation of organisms. For example, exceptions from irreversible somatic differentiation are widespread in plants Lanfear [2018] and are even known in simpler metazoans among cnidarians DuBuc et al. [2020] for which differentiation from vegetative to reproductive functions has been reported. Therefore, the irreversibility of somatic differentiation cannot be taken for granted in the course of the evolution of complex multicellularity.

The majority of the theoretical models addressing the evolution of somatic cells focuses on the evolution of cell specialization, abstracting from the developmental process how germ (reproductive specialists) and soma are produced in the course of the organism growth. For example, a large amount of work focuses on the optimal distribution of reproductive and vegetative functions in the adult organism Michod [2007], Willensdorfer [2009], Rossetti et al. [2010], Rueffler et al. [2012], Ispolatov et al. [2012], Goldsby et al. [2012], Solari et al. [2013], Goldsby et al. [2014], Amado et al. [2018], Tverskoi et al. [2018]. However, these models do not consider the process of organism development. Other work takes the development of an organism into account to some extent: In Gavrilets [2010], the organism development is considered, but the fraction of cells capable to become somatic is fixed and does not evolve. In Erten and Kokko [2020], the strategy of germ-to-soma differentiation is an evolvable trait, but the irreversibility of somatic differentiation is taken for granted. In Rodrigues et al. [2012], irreversible differentiation was found, but both considered cell types pass to the next generation of organisms, such that the irreversible specialists are not truly somatic cells in the sense of evolutionary dead ends. Finally, in Cooper and West [2018] all model ingredients are present: the strategy of cell differentiation is explicitly considered and it is an evolvable trait, also soma and germ cells are considered. However, irreversible somatic differentiation was not observed in that study. Hence, the theoretical understanding of the evolution of irreversibly differentiated somatic cell lines is limited so far.

We developed a theoretical model to investigate conditions for the evolution of the irreversible somatic differentiation, in which vegetative soma-role cells are, in principle, capable to re-differentiate and produce reproductive germ-role cells. In our model, we incorporate factors including (i) costs of cell differentiation, (ii) benefits provided by presence of soma-role cells, (iii) maturity size of the organism. We ask under which circumstances irreversible somatic differentiation is a strategy that can maximize the population growth rate compared to strategies in which differentiation does not occur or somatic differentiation is reversible.

## 2 Model

We consider a large population of clonally developing organisms composed of two types of cells: germ-role and soma-role. Each organism is initiated as a single germ-role cell. In the course of the organism growth, germ-role cells may differentiate to give rise to soma-role cells and vice versa, see Fig. 1A,B. We assume that somatic cells accelerate growth: an organism containing more somatic cells grows faster. After *n* rounds of synchronous cell divisions, the organism reaches its maturity size of 2^*n*^ cells. Immediately upon reaching maturity, the organism reproduces: germ-role cells disperse and each becomes a newborn organism, while all soma-role cells die and are thus lost, see Fig. 1A.

**Figure 1:**
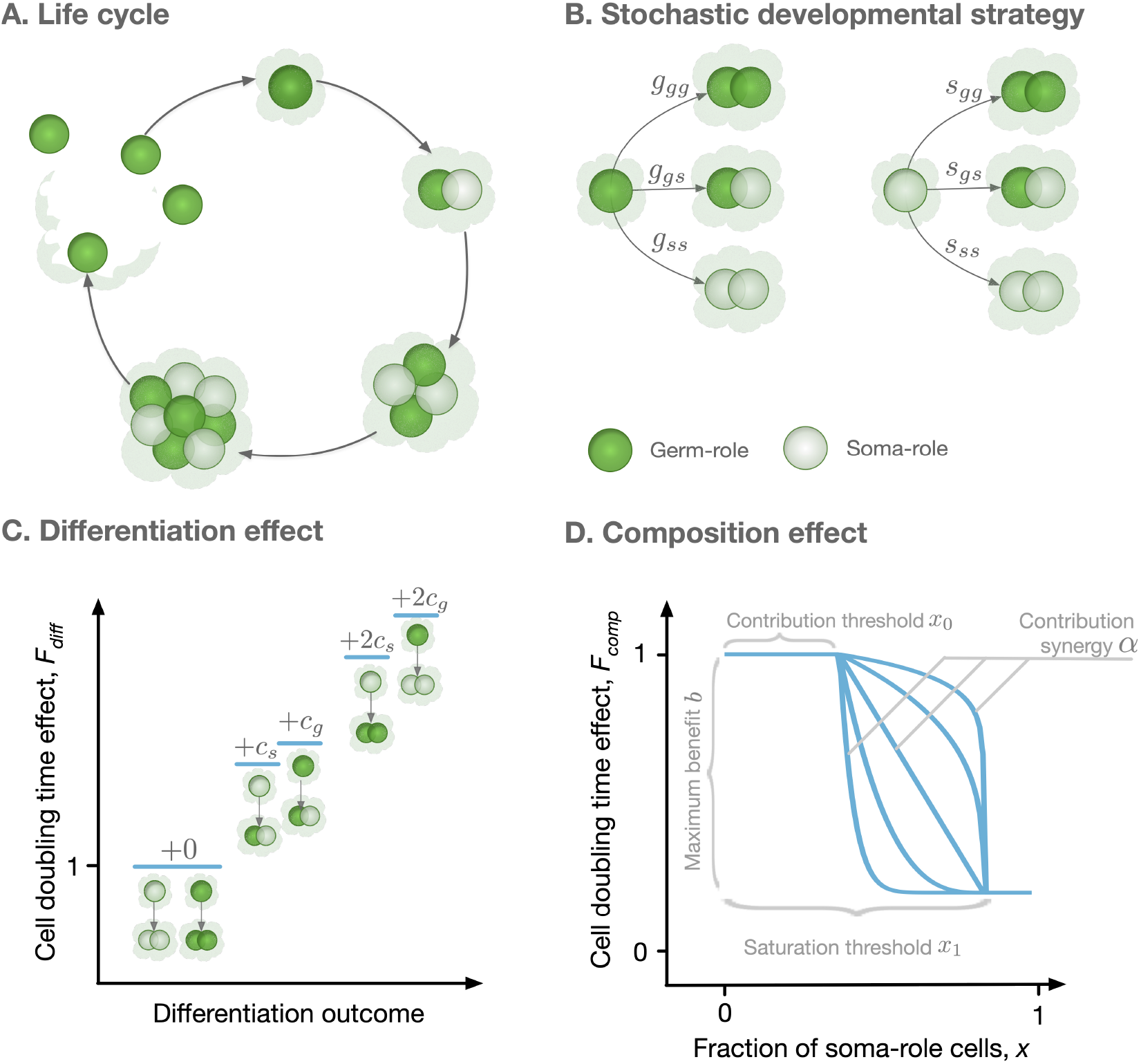
Model overview. **A.** The life cycle of an organism starts with a single germ-role cell. In each round, all cells divide and daughter cells can differentiate into a role different from the maternal cell’s role. When the organism reaches maturity, it reproduces: each germ-role cell becomes a newborn organism and each soma-role cell dies. **B.** Change of cell roles is controlled by a stochastic developmental strategy defined by probabilities of each possible outcomes of a cell division. **C.** Differentiation of cells requires an investment of resources and, thus, slows down the organism growth. Each cell differentiation event incurs a cost (*c_s_* or *c_g_*). The average cost of differentiation contributes increases the cell doubling time in a multiplicative way. **D.** The growth contribution of somatic cells is controlled by a function that decreases the doubling time with the fraction of somatic cells. The form of this function is controlled by four parameters, *x*_0_, *x*_1_, *α*, and *b*.

To investigate the evolution of irreversible somatic differentiation, we consider organisms in which the functional role of the cell (germ-role or soma-role) is not necessarily inherited. When a cell divides, the two daughter cells can change their role, leading to three possible combinations: two germ-role cells, one germ-role cell plus one soma-role cell, or two soma-role cells. We allow all these outcomes to occur with different probabilities, which also depend on the parental type, see Fig 1B. If the parental cell had the germ-role, the probabilities of each outcome are denoted by *g_gg_*, *g_gs_*, and *g_ss_* respectively. If the parental cell had the soma-role, these probabilities are *s_gg_*, *s_gs_*, and *s_ss_*. Altogether, six probabilities define a stochastic developmental strategy *D* = (*g_gg_, g_gs_, g_ss_*; *s_gg_, s_gs_, s_ss_*). In our model, it is the stochastic developmental strategy that is inherited by offspring cells rather than the functional role of the parental cell.

To feature irreversible somatic differentiation (ISD in the following), the developmental strategy must allow germ-role cells to give rise to soma-role cells (*g_gg_* < 1) and must forbid soma-role cells to give rise to germ-role cells (*s_ss_* = 1). All other developmental strategies can be broadly classified into two classes. Reversible somatic differentiation (RSD) describes strategies where cells of both roles can give rise to each other: *g_gg_* < 1 and *s_ss_* < 1. In the strategy with no somatic differentiation (NSD), soma-role cells are not produced in the first place: *g_gg_* = 1, see Table 1.

**Table 1:** Classification of developmental strategies

| Class | Label | $g_{gg}$ | $s_{ss}$ |
| --- | --- | --- | --- |
| Irreversible somatic differentiation | ISD | $< 1$ | $= 1$ |
| Reversible somatic differentiation | RSD | $< 1$ | $< 1$ |
| No somatic differentiation | NSD | $= 1$ | irrelevant |

In our model, evolution is driven by the growth competition between populations executing different developmental strategies. Growth competition will favour developmental strategies that lead to faster growth Pichugin et al. [2017], Gao et al. [2019]. The rate of population growth is determined by the number of offspring produced by an organism (equal to the number of germ-role cells at the end of life cycle) and the time needed for an organism to develop from a single cell to maturity (improved with the number of soma-role cells during the life cycle). The development consists of *n* rounds of simultaneous cell divisions. Consequently, the total development time is a sum of *n* time intervals between cell doubling events. Each cell doubling time *t* is determined by two independent effects: the differentiation effect *F*_diff_ representing costs of changing cell roles Gallon [1992] and the organism composition effect *F*_comp_ representing benefits from having soma-role cells Grosberg and Strathmann [1998, 2007], Shelton et al. [2012], Matt and Umen [2016],

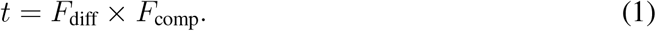

The cell differentiation effect *F*_diff_ represents the costs of cell differentiation. The differentiation of a cell requires efforts to modify epigenetic marks in the genome, recalibration of regulatory networks, synthesis of additional and utilization of no longer necessary proteins. This requires an investment of resources and therefore an additional time to perform cell division. Hence, any cell, which is about to give rise to a cell of a different role, incurs a differentiation costs *c_g_* for germ-to-soma and *c_s_* for soma-to-germ transitions, see Fig. 1C. The resulting effect of differentiation costs is determined as *F*_diff_ = 1 + 〈*c*〉, where 〈*c*〉 is the average differentiation cost among all cells in an organism.

The composition effect profile *F*_comp_(*x*) captures how the cell division time depends on the proportion of soma-role cells *x* present in an organism. In this study, we use a functional form illustrated in Fig. 1D and given by

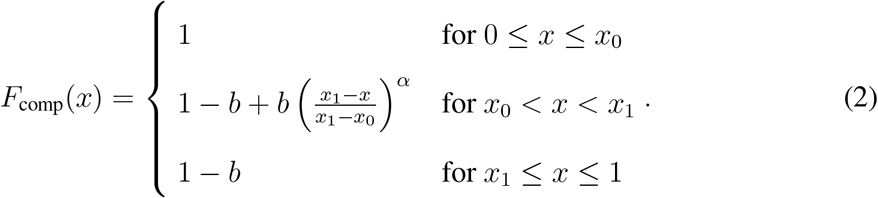

With the functional form (2), soma-role cells can benefit to the organism growth, only if their proportion in the organism exceeds the contribution threshold *x*_0_. Interactions between soma-role cells may lead to the synergistic (soma-role cells work better together than alone), or discounting benefits (soma-role cells work better alone than together) to the organism growth, controlled by the contribution synergy parameter *α*. The maximal achievable reduction in the cell division time is given by the maximal benefit *b*, realized beyond the saturation threshold *x*_1_ of the soma-role cell proportion. A further increase in the proportion of soma-role cells does not provide any additional benefits. With the right combination of parameters, (2) is able to recover various characters of soma-role cells contribution to the organism growth: linear (*x*_0_ = 0, *x*_1_ = 1, *α* = 1), power-law (*x*_0_ = 0, *x*_1_ = 1, *α* ≠ 1), step-functions (*x*_0_ = *x*_1_), and a huge range of other scenarios.

For a given combination of differentiation costs (*c_g_*, *c_s_*) and a composition effect profile (determined by four parameters: *x*_0_, *x*_1_, *b*, and *α*), we screen through a number of stochastic developmental strategies *D* and identify the one providing the largest growth rate to the population. In this study, we searched for those parameters under which ISD strategies lead to the fastest growth and are thus evolutionary optimal, see model details in Appendix A.1.

## 3 Results

### 3.1 For irreversible somatic differentiation to evolve, cell differentiation must be costly

We found that irreversible somatic differentiation (ISD) does not evolve when cell differentiation is not associated with any costs (*c_s_* = *c_g_* = 0), see Fig 2A. This finding comes from the fact that when somatic differentiation is irreversible, the fraction of germ-role cells can only decrease in the course of life cycle. As a result, ISD strategies deal with the tradeoff between producing more soma-role cells at the beginning of the life cycle, and having more germ-role cells by the end of it. On the one hand, ISD strategies which produce a lot of soma-role cells early on, complete the life cycle quickly but preserve only a few germ-role cells by the time of reproduction. On the other hand, ISD strategies which generate a lot of offspring, can deploy only a few soma-role cells at the beginning of it and thus their developmental time is inevitably longer. By contrast, reversible somatic differentiation strategies (RSD) do not experience a similar tradeoff, as germ-role cells can be generated from soma-role cells. As a result, RSD allows higher differentiation rates and can develop a high soma-role cell fraction in the course of the organism growth and at the same time have a large number of germ-role cells by the moment of reproduction. Under costless cell differentiation, for any ISD strategy, we can find an RSD counterpart, which leads to faster growth: the development proceeds faster, while the expected number of produced offspring is the same, see Appendix A.2for details. As a result, costless cell differentiation cannot lead to irreversible somatic differentiation.

**Figure 2:**
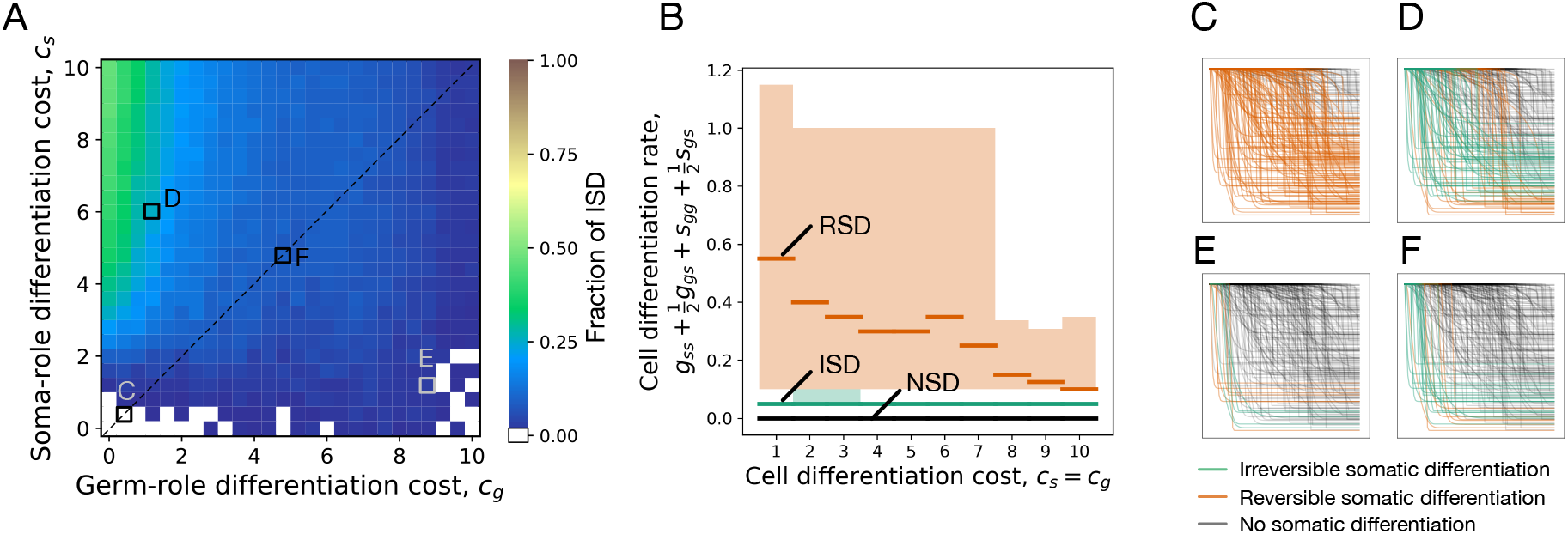
Irreversible soma evolves when cell differentiation is costly. **A.** The fraction of composition effect profiles, (2), promoting ISD as a function of the differentiation costs *c_g_* and *c_g_*. We randomly draw the parameters in (2) to construct 200 random profiles (see Appendix for details). The absence of costs (*c_g_* = *c_s_* = 0) as well as large costs of germ differentiation (large *c_g_*) suppresses the evolution of ISD. Irreversible somatic differentiation is promoted most when the cell differentiation cost is large for soma-role cells (*c_s_*) and small for germ-role cells (*c_g_*). The maturity size used in the calculation is 2^10^ cells. Black dashed lines at panel B indicates the line of equal costs *c_s_* = *c_g_* and squares indicate the costs shown in panels C-F. **B.** Cumulative cell differentiation rate 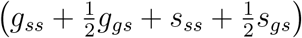 in developmental strategies evolutionarily optimal at various differentiation costs (*c_s_* = *c_g_*), separated by class (ISD, RSD, or NSD). Thick lines represent median values within each class, shaded areas show 90% confidence intervals. For each cost value, 3000 random profiles are used. Evolutionary optimal RSD strategies (orange) have much higher rates of cell differentiation than ISD (green). Consequently, RSD is penalized more under costly differentiation. **C - F.** Shapes of composition effect profiles (compare Fig. 1D) promoting ISD (green lines), RSD (orange lines), and NSD (black lines) developmental strategies at four parameter sets indicated in panel A.

To confirm the reasoning that RSD strategies gain an edge over ISD by having larger differentiation rates, we asked which ISD and RSD strategies become optimal at various cell differentiation costs (*c_s_* = *c_g_*). At each value of costs, we found evolutionarily optimal developmental strategy for 3000 different randomly sampled composition effect profiles *F*_comp_(*x*). We found that evolutionarily optimal RSD strategies feature much larger rates of cell differentiation than evolutionarily optimal ISD strategies, see Fig. 2B. Even at large costs, where frequent differentiation is heavily penalized, the distinction between differentiation rates of ISD and RSD strategies remains apparent.

We screened through a spectrum of germ-to-soma (*c_g_*) and soma-to-germ (*c_s_*) differentiation costs, see Fig 2A. Both differentiation costs punish RSD strategies severely due to their high differentiation rates. By contrast, strategies with irreversible somatic differentiation are insensitive to changes in soma-to-germ differentiation costs *c_s_*, because soma-role cells never give rise to germ-role cells in ISD. Consequently, we observed that ISD is most likely to evolve, when the transition from germ-role to soma-role is cheap (*c_g_* is small) and the reverse transition is expensive (*c_s_* is large), see Fig 2A. In a similar manner, an increase in germ-to-soma differentiation costs (*c_g_*) punishes both RSD and ISD strategies. However, RSD strategies tend to have larger rates of germ-to-soma transitions. Thus, they are punished more than ISD, which leads to the evolution of ISD at small *c_s_* and large *c_g_*. Finally, the NSD strategy does not pay any costs at all, as no cell differentiation occurs. Hence, at very large germ-to-soma differentiation costs (*c_g_* ≈ 10 at Fig. 2A), the NSD strategy outcompetes both reversible and irreversible somatic differentiation, see Appendix A.3for details. For simplicity, hereafter we focus on the case of the equal differentiation costs *c_s_* = *c_g_* = *c* (a black dashed line on Fig 2A).

### 3.2 Evolution of irreversible somatic differentiation is promoted when even a small number of somatic cells provides benefits to the organism

The composition effect profiles *F*_comp_(*x*) that promote the evolution of irreversible somatic differentiation have certain characteristic shapes, see 2C-F. We investigated what kind of composition effect profiles can make irreversible somatic differentiation become an evolutionary optimum. We sampled a number of random composition effect profiles with independently drawn parameter values and found optimal developmental strategies for each profile for a number of differentiation costs (*c*) and maturity size (2^*n*^) values. We took a closer look at the instances of *F*_comp_(*x*) which resulted in irreversible somatic differentiation being evolutionarily optimal.

We found that ISD is only able to evolve when the soma-role cells contribute to the organism cell doubling time even if present in small proportions, see Fig. 3A,B. Analysing parameters of the composition factors promoting ISD, we found that this effect manifests in two patterns. First, the contribution threshold value (*x*_0_) has to be small, see Fig 3D – ISD is promoted when soma-role cells begin to contribute to the organism growth even in low numbers. Second, the contribution synergy was found to be large (*α* > 1) or, alternatively, the saturation threshold (*x*_1_) was small, see Fig 3C.

**Figure 3:**
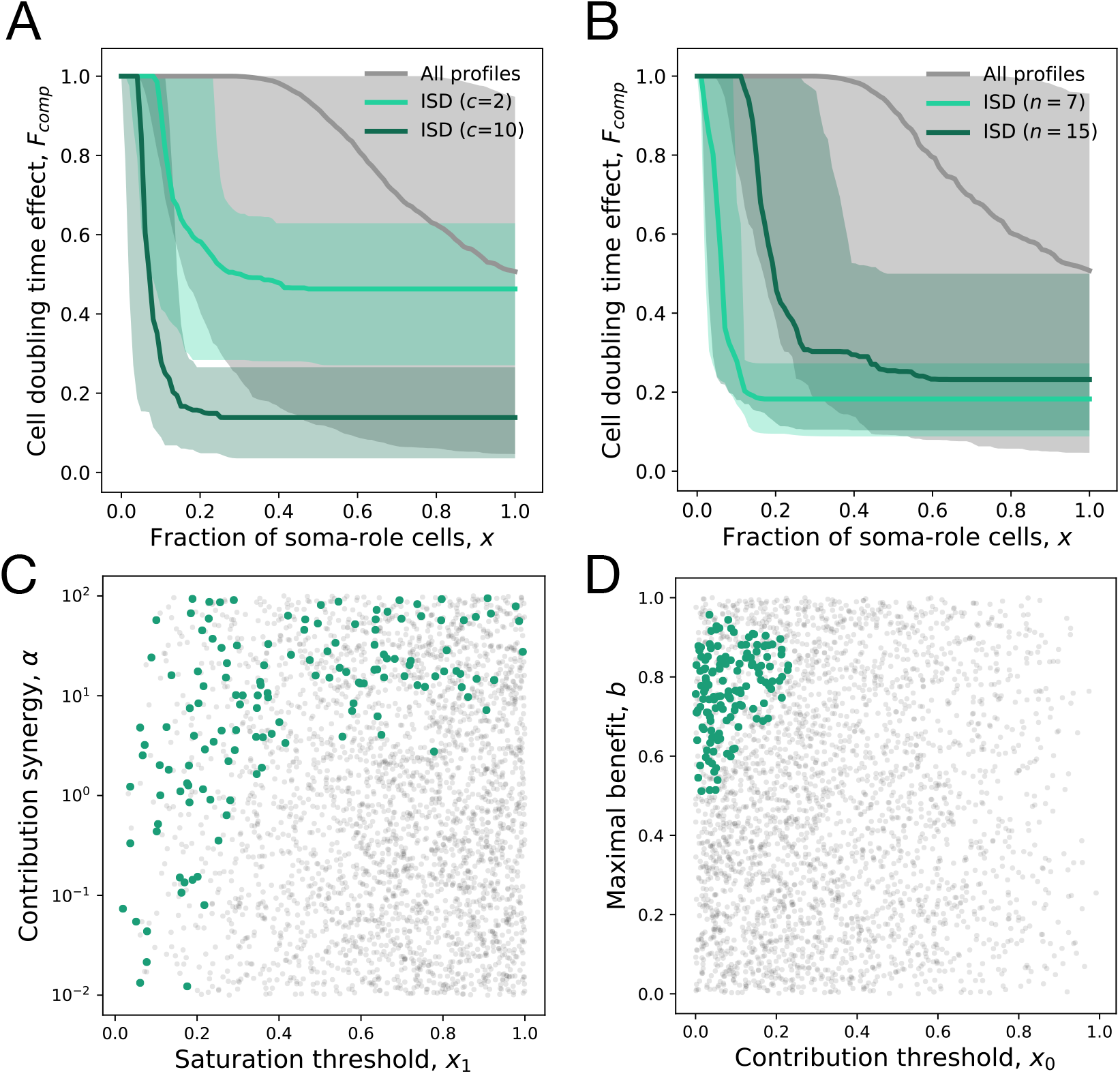
Irreversible soma evolves when substantial benefits arise at small concentrations of soma-role cells. In all panels, the data representing the entire set of composition effect profiles *F*_comp_(*x*) is presented in grey, while the subset promoting ISD is coloured. **A, B.** Median and 90% confidence intervals of composition effect profiles at different differentiation costs (**A**, maturity size *n* = 10) and maturity sizes (**B**, differentiation costs *c* = 5). **C, D.** The set of composition effect profiles in the parameter space. Each point represents a single profile (*c* = 5 and *n* = 10). **C.** The co-distribution of the saturation threshold (*x*_1_) and the contribution synergy (*α*) reveals that either *x*_1_ must be small or *α* must be large. **D.** Co-distribution of the contribution threshold (*x*_0_) and the maximal benefit (*b*) shows that *x*_0_ must be small, while *b* must be large to promote ISD. 3000 profiles are used for panels A, C, D and 1000 profiles for panel B.

Both the contribution threshold *x*_0_ and the contribution synergy *α* control the shape of the composition effect profile at intermediary abundances of soma-role cells. If the contribution synergy *α* exceeds 1, the profile is convex, so the contribution of soma-role cells quickly becomes close to maximum benefit (*b*). A small saturation threshold (*x*_1_) means that the maximal benefit of soma is achieved already at low concentrations of soma-role cells (and then the shape of composition effect profile between two close thresholds has no significance). Together, these patterns give an evidence that the most crucial factor promoting irreversible somatic differentiation is the effectiveness of soma-role cells at small numbers, see Appendix A.4for more detailed data presentation.

The reason behind these patterns is a slower accumulation of soma-role cells under irreversible somatic differentiation, comparing to RSD strategies, see Appendix A.2. Thus, with ISD, an organism spends a significant amount of time having only a few soma role cells. Hence, ISD strategy can only be evolutionarily successful, if these few soma-role cells have a notable contribution to the organism growth time.

We also found that profiles featuring ISD do not possess neither extremely large, nor extremely small maximal benefit values *b*, see Fig. 3D. When the maximal benefit is too small, the cell differentiation just does not provide enough benefits to be selected for and the evolutionarily optimal strategy is NSD. In the opposite case, when the maximal benefit is very close to one, the cell doubling time approaches zero, see (2). Then, the benefits of having many soma-role cells outweighs the costs of differentiation and the optimal strategy is RSD, see Appendix A.4.

### 3.3 For irreversible somatic differentiation to evolve, the organism size must be large enough

By screening through the maturity size (2^*n*^) and differentiation costs (*c*), we found that the evolution of irreversible somatic differentiation is heavily suppressed at small maturity sizes, Fig 4A. For *c_s_* = *c_g_*, the minimal maturity size allowing irreversible somatic differentiation to evolve is 2^*n*^ = 64 cells. At the same time, organisms performing just a few more rounds of cell divisions are able to evolve ISD at a wide range of cell differentiation costs, see also Appendix A.5. This indicates that the evolution of irreversible somatic differentiation is strongly tied to the size of the organism.

**Figure 4:**
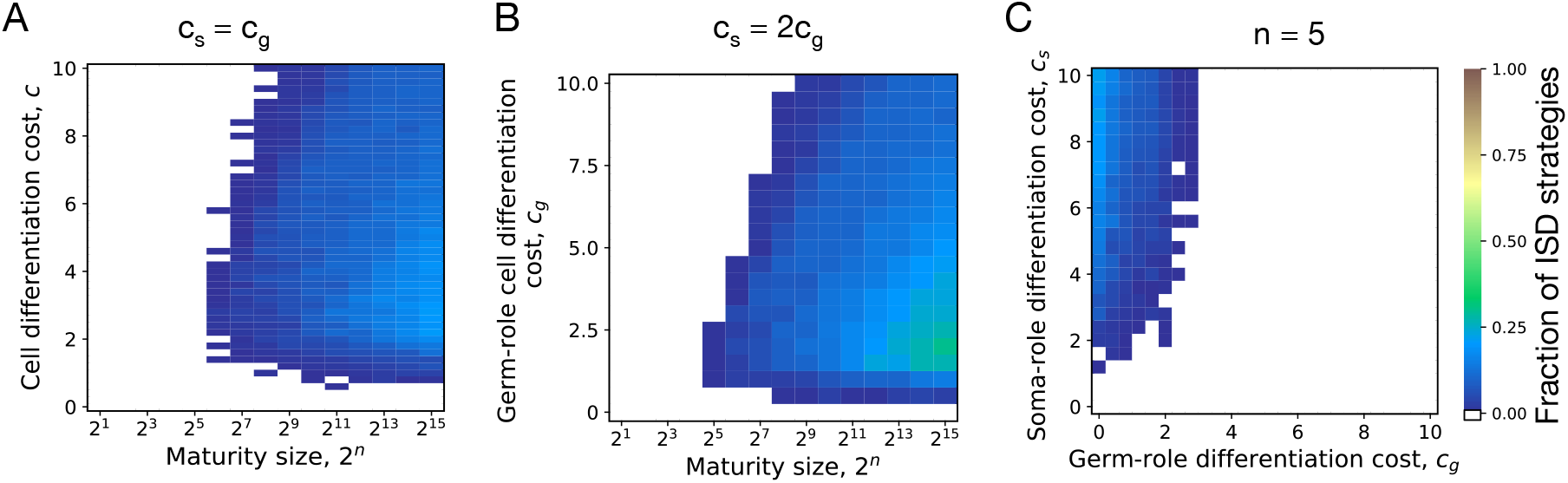
Irreversible soma can evolve if organism grows to a large enough size in the course of its life cycle. **A.** The fraction of composition effect profiles promoting ISD at various cell differentiation costs (*c* = *c_s_* = *c_g_*) and maturity sizes (2^*n*^). ISD strategies were only found for maturity size 2^6^ = 64 cells and larger. **B.** The fraction of composition effect profiles promoting ISD at unequal differentiation costs *c_s_* = 2*c_g_*. A rare occurrences of ISD (∼ 1%) was detected at the maturity size 2^5^ = 32 cells in a narrow range of cell differentiation costs but not at the smaller sizes. **C.** The range of cell differentiation costs promoting ISD at at the maturity size 2^5^ = 32 cells. For ISD strategies to evolve at such a small size, the differentiation from soma-role to germ-role must be much more costly than the opposite transition (*c_s_* ≫ *c_g_*).

Evolution of ISD at sizes smaller than 64 cells is possible for *c_s_ > c_g_*. For instance, at *c_s_* = 2*c_g_* some ISD strategies were found to be optimal at the maturity size 2^5^ = 32 cells, Fig 4B. However, ISD strategies were found in a narrow range of cell differentiation costs and the fraction of composition effect profiles that allow evolution of ISD there was quite low – about 1%. The evolution of ISD at such small maturity sizes becomes likely only at extremely unequal costs of transition between germ and some roles *c_s_* ≫ *c_g_*, see Fig 4C. Hence, for irreversible somatic differentiation to evolve, the organism size should exceed a threshold of roughly 64 cells.

## 4 Discussion

The vast majority of cells in a body of any multicellular being contains enough genetic information to build an entire new organism. However, in a typical metazoan species, very few cells actually participate in the organism reproduction – only a limited number of germ cells are capable to do it. The other cells, called somatic cells, perform vegetative functions but do not try to form an offspring organism – somatic differentiation is irreversible. We asked for the reason for the success of such a specific mode of organism development. We theoretically investigated the evolution of irreversible somatic differentiation with a model of clonally developing organisms taking into account benefits provided by soma-role cells, costs arising from cell differentiation, and the effect of the raw organism size.

While our model can capture some key features of these biological systems, it remains of course an abstraction. We assumed that populations go into an exponential growth phase – competition for space or nutrients could lead to selection of other strategies instead. Additional features such as trade-offs in growth at different colony sizes lead to further complications. Nevertheless, our model allows to start to look into the basic features of nascent life cycles at the edge of the division of labour in multicellular colonies.

Our key findings are:

- The evolution of irreversible somatic differentiation is inseparable from cell differentiation being costly.
- For irreversible somatic differentiation to evolve, somatic cells should be able to contribute to the organism performance already when their numbers are small.
- Only large enough organisms tend to develop irreversible somatic differentiation.

According to our results, cell differentiation costs are essential for the emergence of irreversible somatic differentiation, see Fig. 2A. For cells in a multicellular organism, differentiation costs arise from the material needs, energy, and time it takes to produce components necessary for the performance of the differentiated cell, which were absent in the parent cell. For instance, in filamentous cyanobacteria nitrogen-fixating heterocysts develop much thicker cell wall than parent photosynthetic cells had. Also, reports indicate between 23% Ow et al. [2008] and 74% Sandh et al. [2014] of the proteome changes its abundance in heterocysts compared against photosynthetic cells. Similarly, the changes in the protein composition in the course of cell differentiation was found during the development of stalk and fruiting bodies of *Dictyostelium discoideum* Bakthavatsalam and Gomer [2010], Czarna et al. [2010] Our model demonstrates that irreversible somatic differentiation is more likely to evolve when a few soma-role cells are able to provide a substantial benefit to the organism, see Fig. 3. Several patterns of how the benefit provided by somatic cells changes with their numbers have been previously considered in the literature. However, the range of studied examples was restricted to concave or convex shapes Michod [2007], Willensdorfer [2009], Rossetti et al. [2010], Cooper and West [2018]. In this paper, we went beyond these shapes and additionally considered lower (*x*_0_) and upper (*x*_1_) thresholds for the somatic cells contribution (our model recovers the previous approaches for *x*_0_ = 0 and *x*_1_ = 1). While our findings are in a qualitative agreement with these past results – the profiles promoting irreversible somatic differentiation appear convex, see Fig. 3A,B – our model indicates that the crucial component here is the large benefits provided by small numbers of soma-role cells, rather than overall convexity of the profile. For example, with sufficiently small *x*_1_, the non-constant section of the composition effect profile (where the fraction of soma-role cells is between *x*_0_ and *x*_1_, see Fig. 1D) can be concave (*α* < 1, see Fig. 3C) and still promote irreversible somatic differentiation. *Volvocales* algae demonstrate that a significant contribution by small numbers of somatic cells might indeed be found in a natural population: In *Eudorina illinoiensis* – one of the simplest species demonstrating the first signs of reproductive division of labour, only four out of thirty-two cells are vegetative Sambamurty [2005] (soma-role in our terms). This species has developed some reproductive division of labour and a fraction of only 1/8 of vegetative cells is sufficient for colony success. Thus, it seems possible that highly-efficient soma-role cells open the way to the evolution of irreversible somatic differentiation.

Our model shows that irreversible somatic differentiation can only emerge in relatively large organisms, see Fig 4A. The maturity size plays an important role in an organism’s life cycle Amado et al. [2018], Erten and Kokko [2020]: Large organisms have potential advantages to optimize themselves in multiple ways, such as to improve growth efficiency Waters et al. [2010], to avoid predators Matz and Kjelleberg [2005], Fisher et al. [2016], Hiltunen and Becks [2014] to increase problem-solving efficiency Morand-Ferron and Quinn [2011], and to exploit the division of labour in organisms Carroll [2001], Matt and Umen [2016]. Moreover, the maximum size has been related to the reproduction of the organism from the onset of multicellularity in Earth’s history Ratcliff et al. [2012]. Our results suggest that the smallest organism able to evolve irreversible somatic differentiation should typically be about 32 − 64 cells (unless the cost of soma-to-germ differentiation is extremely large and the cost of the reverse is low). This is in line with the pattern of development observed in *Volvocales* green algae. In *Volvocales*, cells are unable to move (vegetative function) and divide (reproductive function) simultaneously, as a unique set of centrioles are involved in both tasks Wynne and Bold [1985], Koufopanou [1994]. *Chlamydomonas reinhardtii* (unicellular) and *Gonium pectorale* (small colonies up to 16 cells) perform these tasks at different times. They move towards the top layers of water during the day to get more sunlight. At night, however, these species perform cell division and/or colony reproduction, slowly sinking down in the process. However, among larger *Volvocales*, a division of labour begins to develop. In *Eudorina elegans* colonies, containing 16 - 32 cells, a few cells at the pole have their chances to give rise to an offspring colony reduced Marchant [1977], Hallmann [2011]. In *Pleodorina californica*, half of the 128-celled colony is formed of smaller cells, which are totally dedicated to the colony movement and die at the end of colony life cycle Kikuchi [1978], Hallmann [2011]. In *Volvox carteri*, most of a 10000-cell colony is formed by somatic cells, which die upon the release of offspring groups Hallmann [2011].

Our study originated from curiosity about driving factors in the evolution of irreversible somatic differentiation: Why does the green algae *Volvox* from the kingdom Plantae shed most of its biomass in a single act of reproduction? And why, in another kingdom, Animalia, in most of the species the majority of body cells is outright forbidden to contribute to the next generation? Our results show which factors makes a difference between the evolution of an irreversible somatic differentiation and other strategies of development. One of these factors, the maturity size, is known in the context of the evolution of reproductive division of labour Kirk [2005]. Another factor, the costs of cell differentiation, is, in general, discussed in a greater biological scope but is hardly acknowledged as a factor contributing to the evolution of organism development. Finally, the early contribution of soma-role cells to the organism growth, even if they are small in numbers, is an unexpected outcome of our investigation, overlooked so far as well. Despite the simplistic nature of our model (we did not aim to model any specific organism), all our results find a confirmation among the *Volvocales* clade. Hence, we expect that the findings of this study reveal general properties of the evolution of irreversible somatic differentiation, independently of the clade where it evolves.

## A Appendix

## A.1 Search for the evolutionarily optimal developmental program

## A.1.1 Finding the population growth rate for a given developmental program

In [Gao et al., 2019], we have shown that a population of organisms, which begin their life cycle from the same state but have a stochastic development, eventually grows exponentially with the rate *λ* given by the solution of

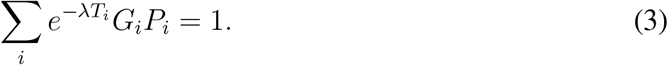

Here, *i* is the developmental trajectory – in our case, the specific combination of all cell division outcomes; *P_i_* is the probability that an organism development will follow the trajectory *i*; *T_i_* is the time necessary to complete the trajectory *i* – from a single cell to the maturity size of 2^*n*^ cells; *G_i_* is the number of offspring organisms produced at the end of developmental trajectory *i*, equal to the number of germ-role cells at the moment of maturity.

In order to find the population growth rate, we need to know *G_i_*, *T_i_*, and *P_i_* (how many offspring are produced, how long did it take to mature, and how likely is this developmental trajectory, respectively). The complete set of developmental trajectories is huge as it scales exponentially with the number of divisions *n*.

In our study, for each developmental strategy, we sampled *M* = 300 developmental trajectories at random. To get each trajectory, we simulated the growth of the single organism according to the rules of our model. For each trajectory, the developmental time *T_i_* was computed as a sum of cell doubling times at each of the *n* synchronous cell divisions, the number of offspring *G_i_* was given by the count of germ-role cells at the end of development. The resulting ensemble of trajectories (with *P_i_* = 1*/M*) was plugged into (3) to compute the population growth rate *λ*.

## A.1.2 Finding the developmental program with the largest population growth rate

We assume that evolution occurs by growth competition between populations executing different developmental strategies. These strategies, which provide larger population growth rate will outgrow others. To find evolutionarily optimal strategies under given conditions, we screened through a large set of developmental strategies and identified the one with the maximal population growth rate *λ*. Since the probabilities of cell division outcomes sum into one (*g_gg_* +*g_gs_*+*g_ss_* = 1 and *s_gg_* +*s_gs_*+*s_ss_* = 1), these probabilities can be represented as a point on two simplexes, one for the division of germ-role cells, and one for the division of soma-role cells. Consequently, we choose the set of developmental strategies as a Cartesian product of two triangular lattices - one for division probabilities of germ-role cells (*g_gg_, g_gs_, g_ss_*) and one for soma-role cells (*s_gg_, s_gs_, s_ss_*). The lattice space was set to 0.1, so each of two independent lattices contained 11 × 12/2 = 66 nodes, and the whole set of developmental strategies comprised 66 × 66 = 4356 different strategies. For each of these strategies, the population growth rate *λ* was calculated and the strategy with the largest growth rate was identified as evolutionarily optimal.

In our investigation, parameters such as differentiation costs (*c_s_*, *c_g_*) and maturity size (2^*n*^) were used as control parameters. In other words, we either fix them at the specific values, or screened through a range of values to obtain a map (see Figs. 2 and 3 in the main text). However, the parameters controlled the shape of composition effect profile (*x*_0_, *x*_1_, *α*, and *b*) were treated differently. For each combination of control parameters, we randomly sampled a number (between 200 and 3000) of combinations of these parameters. The thresholds (0 ≤ *x*_0_ ≤ *x*_1_ ≤ 1) were sampled as a pair of independent distributed random values from the uniform distribution *U* (0, 1). The contribution threshold *x*_0_ was set to the minimum of the pair, and the saturation threshold *x*_1_ was set to the maximum. The contribution synergy (*α* > 0) corresponds to the concave shape of the profile at *α* < 1 and to the convex shape at *α* > 1. Therefore, log_10_(*α*) was sampled from the uniform distribution *U* (−2, +2), so the profile has an equal probability to demonstrate concave and convex shape. Finally, the maximum benefit (0 ≤ *b* < 1) was sampled from a uniform distribution, *U* (0, 1). For each tested combination of control parameters, we found the optimal developmental strategy for every sampled profile. We then classified these as irreversible somatic differentiation (ISD), reversible somatic differentiation (RSD), or no somatic differentiation (NSD).

## A.2 Under costless cell differentiation, irreversible soma strategy cannot be evolutionarily optimal

In this section, we will show that an ISD strategy can never be an evolutionary optimum without cell differentiation being costly. To do that, we first consider the deterministic dynamics of the expected composition of the organism. Then, for an arbitrary ISD strategy, we identify a more advantageous RSD strategy which gives the same organism composition at the end of life cycle but higher number of soma-role cells during the life cycle.

In our model, the composition of the organism is governed by the stochastic developmental strategy and differs between different organisms. Here, as a proxy for this complex stochastic dynamics, we consider the mathematical expectation of the composition. Assume that after *t* ≥ 0 cell divisions the fraction of soma-role cells is *s*(*t*) and the fraction of germ-role cells is *g*(*t*) = 1 − *s*(*t*). Then, the expected fractions of cells of the two types after the next cell division is

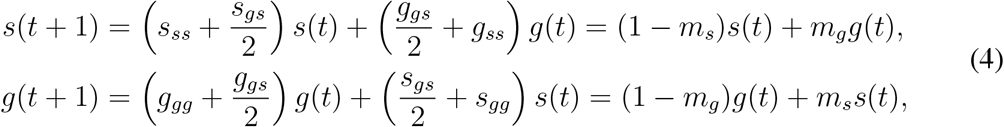

where we introduced 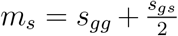 and 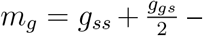 the probabilities that the offspring of a cell will have a different role. Naturally, for irreversible somatic differentiation (ISD) *m_s_* = 0 and *m_g_* > 0, for NSD strategies *m_g_* = 0 and *m_s_* being irrelevant, while the reversible differentiation (RSD) class covers the rest. (4) can be written in matrix form

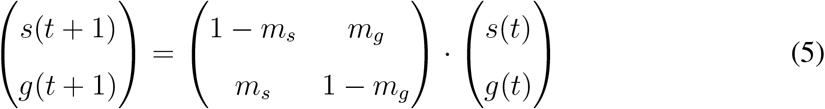

A newborn organism contains a single germ-role cell (*s*(0) = 0, *g*(0) = 1), therefore, the expected composition of an organism after *i* divisions is

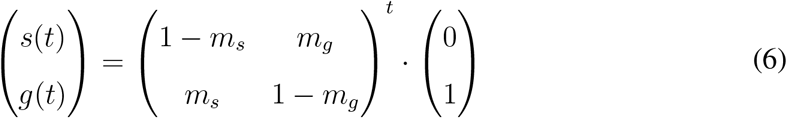

The matrix has two eigenvalues: 1 and 1 − *m_g_* − *m_s_*, with associated right eigenvectors (*m_g_, m_s_*)^*T*^ and (1, −1)^*T*^, respectively. Hence, the expected composition after *t* divisions can be obtained in the explicit form

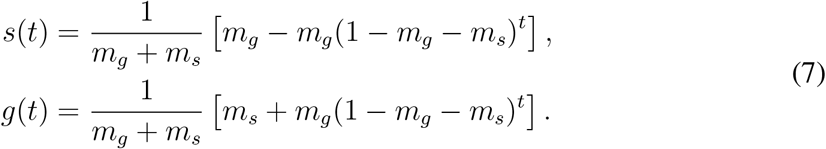

For an arbitrary irreversible somatic differentiation strategy *D*, *m_s_* = 0, the expected number of soma-role cells changes as

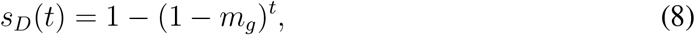

which is a monotonically increasing function of the number of cell divisions *t*, see the green line in Fig. 5. In the life cycle involving *n* cell divisions, the fraction of soma-role cells at the end of life cycle is *s_D_*(*n*) = 1 − (1 − *m_g_*)^*n*^.

**Figure 5:**
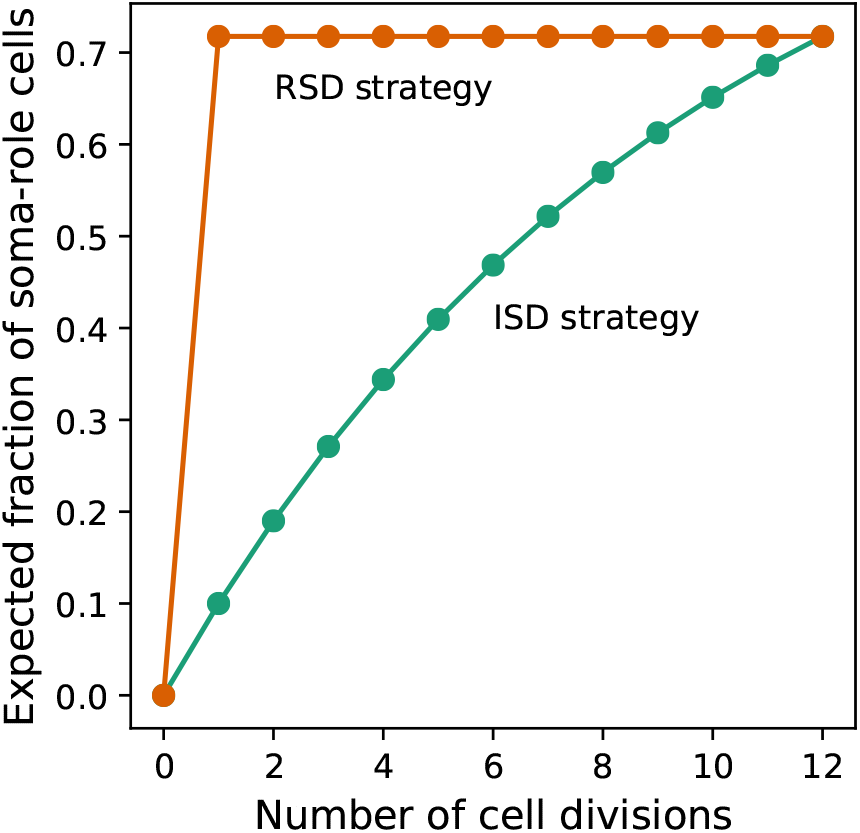
Under costless differentiation, for any irreversible somatic differentiation strategy, exists a reversible somatic differentiation strategy dominating it. The green curve shows the dynamics of the expected fraction of soma-role cells in an organism using an ISD developmental strategy (*m_g_* = 0.1, *m_s_* = 0.0, *n* = 12). The orange curve shows the dynamics of the expected fraction of soma-role cells in an organism using the specific RSD developmental strategy 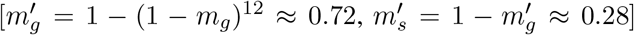. In this strategy, the number of offspring produced at the end of the life cycle is the same as in the considered ISD strategy. At the same time, the fraction of soma-role cells during the life cycle is larger. Therefore, under costless differentiation, the presented RSD strategy is more effective than the considered ISD strategy.

Now, we consider another developmental strategy *D′* with reversible somatic differentiation in which 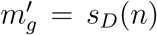 and 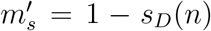. Using 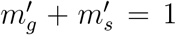 in (7), it can be shown that the expected fraction of soma-role cells in *D′* after the very first cell division is exactly *s_D_*(*n*) and stays constant thereafter, see the orange line in Fig. 5. Thus, the number of offspring produced is the same for both development strategies.

If cell differentiation is costless (*d_s_* = *d_g_* = 0), then the cell doubling time depends only on the fraction of soma-role cells. As all soma-role cells are then present already after the first cell division, organisms following the RSD strategy *D′* will grow faster than organisms using the ISD strategy *D* at any stage of organism development, independently of the choice of the composition effect profile (*F*_comp_). At the end of the life cycle, both strategies have the same expected number of offspring. Therefore, under costless cell differentiation, for any ISD strategy, we can find a RSD strategy that leads to a larger population growth rate.

## A.3 Conditions promoting the evolution of ISD, RSD, and NSD strategies

**Figure 6:**
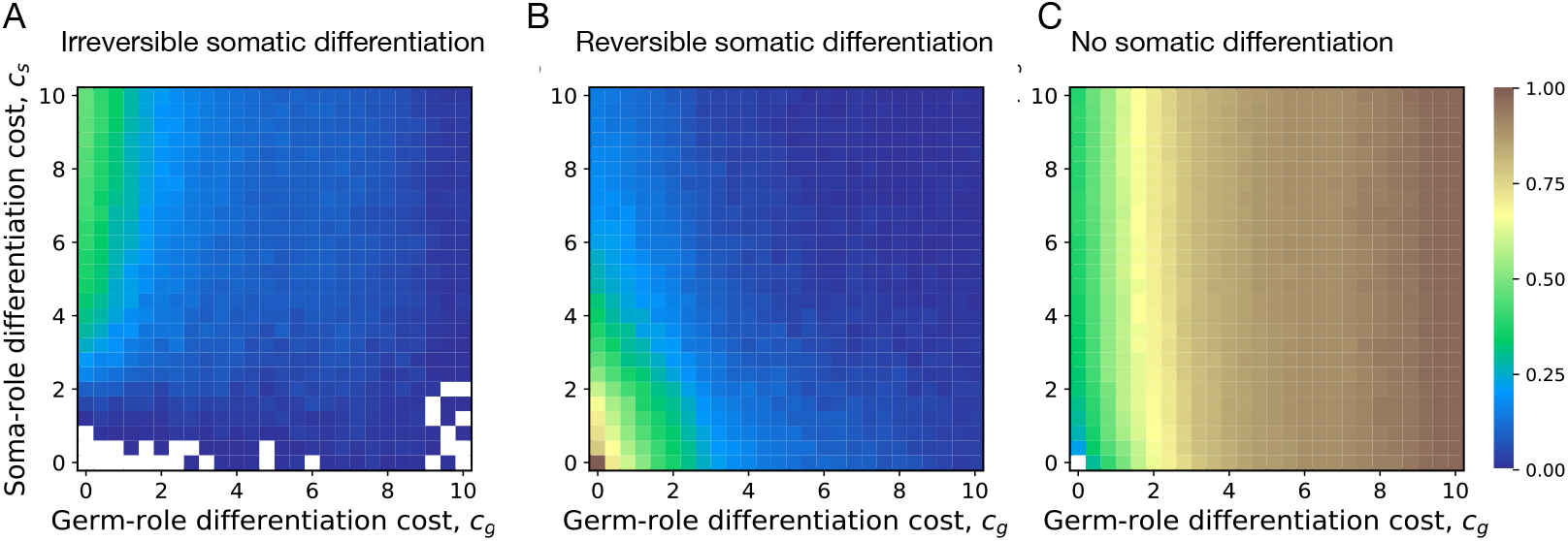
Impact of cell differentiation costs on the evolution of development strategies. The fractions of 200 random composition effect profiles promoting ISD (**A**), RSD (**B**), and NSD (**C**) strategies at various cell differentiation costs (*c_s_*, *c_g_*). In the absence of costs (*c_g_* = *c_s_* = 0), only RSD strategies were observed. RSD strategies are prevalent at smaller cell differentiation costs. NSD strategies are the most abundant at large costs for germ-role cells (*c_g_*). ISD strategies are the most abundant at large costs for soma-role cells (*c_s_*). The maturity size used in the calculation is 2^10^ cells.

**Figure 7:**
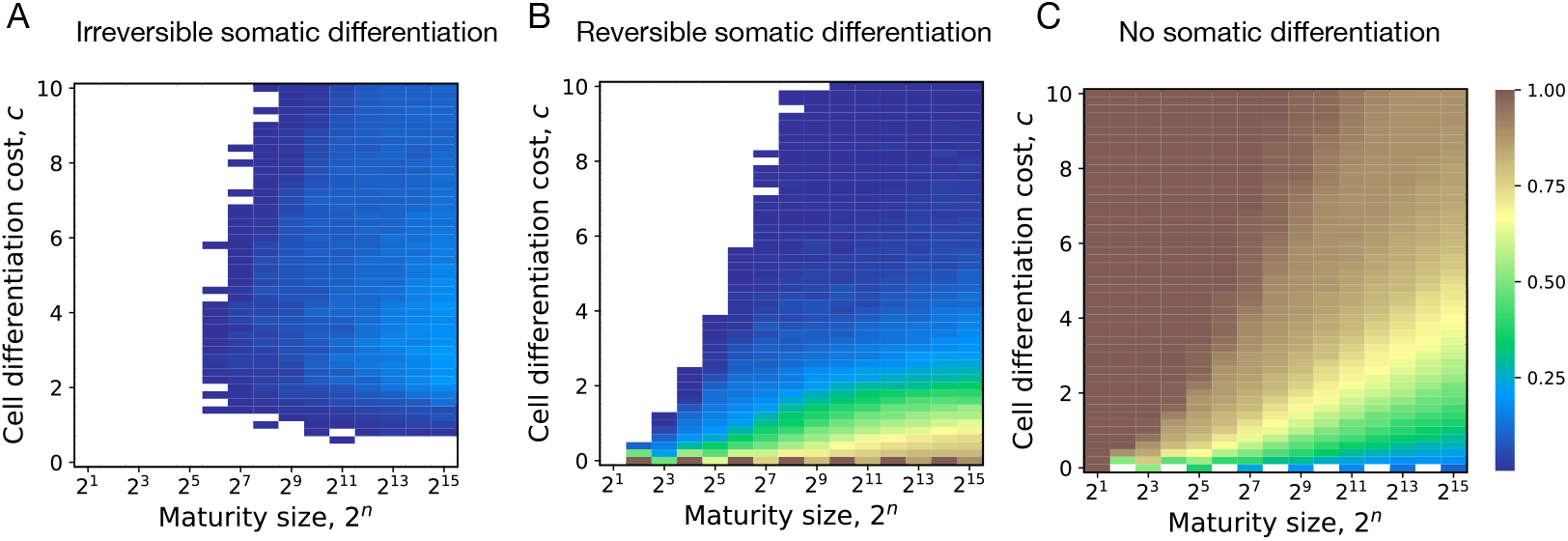
Impact of maturity size on the evolution of development strategies. The fractions of 200 random composition effect profiles promoting ISD (**A**), RSD (**B**), and NSD (**C**) strategies at various cell differentiation costs (*c* = *c_s_* = *c_g_*) and maturity size 2^*n*^. ISD strategies are most abundant at larg maturity sizes and intermediary cell differentiation costs. RSD strategies are most abundant at small cell differentiation costs. NSD strategies are most abundant at small maturity sizes and cell differentiation costs.

## A.4 Parameters of composition effect profiles promoting ISD, RSD, and NSD strategies

**Figure 8:**
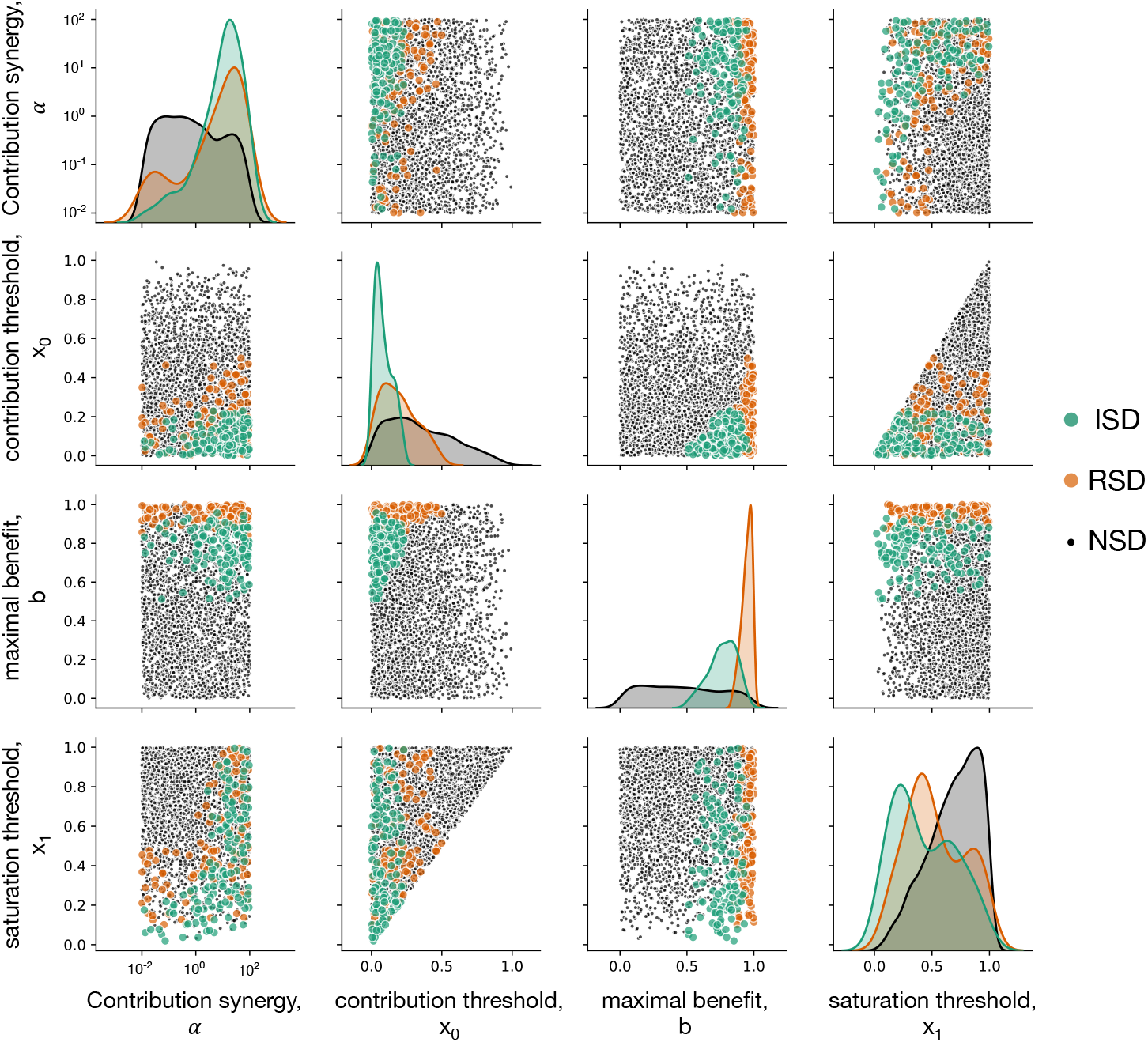
Impact of composition effect parameters on the evolution of development strategies. Each diagonal panel represents individual distribution of each of four parameters among composition effect profiles promoting ISD (green), RSD (orange), and ISD (black) strategies. Each non-diagonal panel represents a pairwise co-distribution of these parameters. ISD strategies are promoted at small contribution thresholds *x*_0_ and for large maximal benefit *b*. Also, either the contribution synergy *α* must be large, or the saturation threshold *x*_1_ should be small - see main text for detailed discussion. RSD strategies require very large *b* - there the benefits of having a large number of soma-role cells outweighs costs paid by frequent differentiation. Due to the fast accumulation of soma-role cells, RSD strategies tolerate larger *x*_0_ than ISD. RSD exhibit the same restrictions with respect to *x*_1_ as ISD and are insensitive to *α*. For this figure, 3000 composition effect profiles were investigated with costs *c* = *c_s_* = *c_g_* = 5 and *n* = 10.

## A.5 Evolution of irreversible somatic differentiation under various maturity sizes and unequal cell differentiation costs

**Figure 9:**
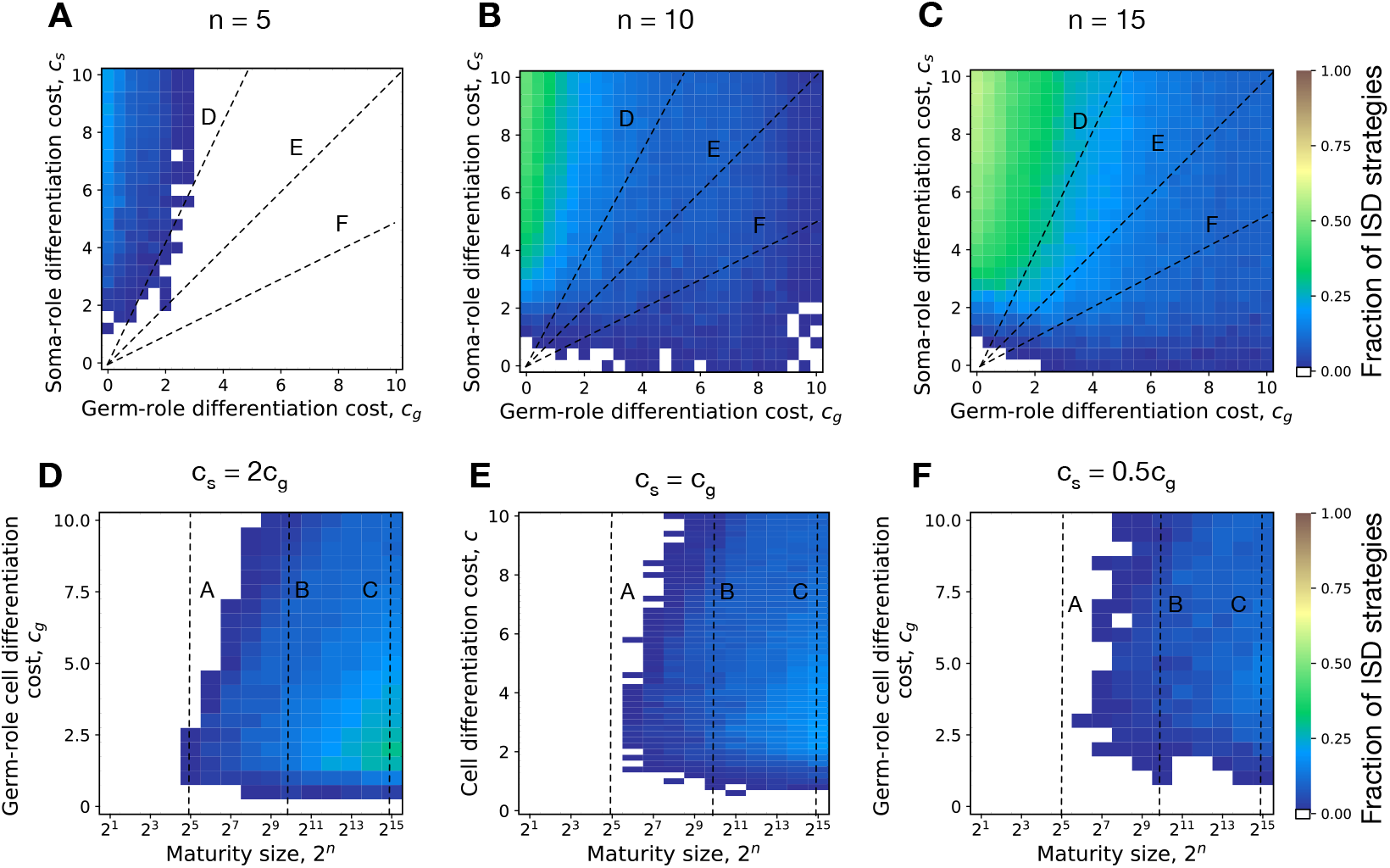
Evolution of irreversible somatic differentiation at unequal cell differentiation costs. **A-C.** The fraction of 200 random composition effect profiles promoting ISD at various cell differentiation costs (*c_s_*, *c_g_*) at fixed maturity size *n* = 5 (panel A), 10 (B), and 15 (C). Larger maturity sizes promote the evolution of ISD across all cell differentiation costs. **D-F.** The fraction of composition effect profiles promoting ISD at unequal cell differentiation costs *c_s_/c_g_* = 2 (panel D), *c_s_/c_g_* = 1 (E), and *c_s_/c_g_* = 0.5 (F). Even with unequal differentiation costs, the minimal maturity size allowing the evolution of ISD stays roughly the same — 2^5^ − 2^6^ cells. Dashed lines indicate overlap between panels.

## Notes

### Competing Interest Statement

The authors have declared no competing interest.

